# Information use shapes range expansion dynamics into environmental gradients

**DOI:** 10.1101/056002

**Authors:** Emanuel A. Fronhofer, Nicolai Nitsche, Florian Altermatt

## Abstract

Globally, geographic distributions of species are dynamic and strongly influenced by dispersal. Vice versa, range dynamics feed back and may select for increased dispersal. This interplay almost universally happens across environmental gradients which can directly impact the fitness of organisms, but also provide individuals with information on the environmental changes. However, the organisms’; ability to subsequently adjust dispersal decisions plastically has been largely ignored and the (macro)ecological consequences remain unclear. Using modeling and controlled experiments in replicated microcosm landscapes, we show that information on environmental gradients severely impacted range dynamics and inverted the spatial distribution of population densities in comparison to controls where this information was not provided. Additionally, information use prevented evolutionary changes in dispersal and an acceleration of range expansions. We demonstrate the strong impact of informed dispersal and subsequent behavioral changes on range dynamics in environmental gradients and spatial dynamics in general.

## Introduction

The capacity of organisms to spread in space and to expand their range into new habitat is crucial for their long-term fitness, especially in the context of current global environmental and climatic changes (Hill *et al.*, 1999; Parmesan *et al.*, 1999; Parmesan & Yohe, 2003; Kelly & Goulden, 2008). The fundamental and applied relevance of range expansions and biological invasions resulted in extensive theoretical work predicting range dynamics (Hastings *et al.*, 2005; Holt *et al.*, 2005; Burton *et al.*, 2010; Dytham, 2009; Holt & Barfield, 2011; Perkins *et al.*, 2013). To date, however, our empirical understanding of range dynamics is mostly based on case studies of range shifts and invasions with little experimental validation or manipulation. The few studies that experimentally track replicated range expansions are either limited by the short time frames considered (Melbourne & Hastings, 2009; Giometto *et al.*, 2014), preventing potentially important evolutionary changes to occur, or by the unrealistic assumption that range expansions occur into uniform habitat (Fronhofer & Altermatt, 2015).

Realistically, all range expansions are limited by the heterogeneity of landscapes and the universally present gradients in environmental conditions, such as temperature or humidity. While the importance of gradients as such for species ranges has been explored previously (Kubisch *et al.*, 2010, 2014; Louthan *et al.*, 2015) these works consistently ignore that environmental gradients always have a two-fold effect on organisms: 1) Gradients have a direct, fitness-relevant effect due to the mismatch between local conditions and the individuals’; environmental optimum. 2) Furthermore, gradients have indirect effects mediated by information on changing local conditions provided to spreading organisms, which subsequently may lead to plastic changes in dispersal and altered range dynamics. While the relevance of information use for making dispersal decisions and subsequent consequences for spatial dynamics has been recognized in general (Clobert *et al.*, 2009), the consequences of informed dispersal for macroecological dynamics, such as species range shifts, remains under-appreciated.

Here, we theoretically and experimentally test the role of environmental gradients for the dynamics of range expansions taking into account the two-fold effect of environmental gradients discussed above. We use an individual-based model to predict ecological and evolutionary dynamics in three range expansion scenarios: Firstly (“control”), we model the range expansion of individuals into a previously empty linear landscape of interconnected patches. Secondly, we include a scenario analogous to the first, but where the landscape is characterized by a linear gradient of increasing local mortality that affects the spreading organisms’; fitness without providing information on the spatial change in mortality (“gradient”). Finally, we contrast these two scenarios with a range expansion into a mortality gradient that provides information on the changes in mortality and individuals use this information to make optimal dispersal decision plastically (“gradient & information”).

We tested our theoretical predictions using experimental evolution and replicated linear microcosm landscapes, which were invaded by the ciliate model organism *Tetrahymena pyriformis* (Altermatt *et al.*, 2015). The landscapes allowed for active dispersal and included the three scenarios detailed above: control, gradient as well as gradient & information.

We predict that range expansions in the control scenario lead to the evolution of increased dispersal at the range front (Phillips *et al.*, 2006; Fronhofer & Altermatt, 2015). The mortality gradient in the second scenario should lead to a reduction in range expansion speed and, ultimately, to the establishment of a stable range border due to the ecological effect of the mortality gradient on local population dynamics and the evolutionary effect selecting against dispersal (Kubisch *et al.*, 2014). Finally, the availability of information should provide organisms with the opportunity to make informed and plastic dispersal decisions and thereby not to disperse into areas characterized by high local mortalities.

## Materials and Methods

### Numerical analyses

#### General overview

We developed a stochastic, individual-based simulation model (Burton *et al.*, 2010; Kubisch *et al.*, 2014; Fronhofer & Altermatt, 2015) that tracks, firstly, ecological dynamics, such as spatial spread in a linear landscape, population densities as well as dispersal events, and, secondly, evolutionary changes, more specifically the evolution of dispersal and the concurrent evolution of reproductive and competitive ability. In each replicate linear landscape, populations are initialized at one end of the landscape and individuals may subsequently spread following a stepping stone model (nearest neighbor dispersal).

We assume local competition for resources and, for simplicity, non-overlapping generations. As a result of standing genetic variation present in the beginning and of subsequent mutations, the distribution of traits in a population may shift, leading to evolutionary changes in dispersal. Since it is well known that dispersal is costly (Bonte *et al.*, 2012), we assume that more dispersive individuals reproduce less due to their investment of energy into dispersal (Fronhofer & Altermatt, 2015) (dispersal-fecundity tradeoff; Eq. 4). Furthermore, reproduction and competitive ability are positively correlated (Eq. 3) due to underlying consumer-resource dynamics (Matessi & Gatto, 1984) (detailed derivation in the Electronic Supplementary Material).

In addition to a control scenario (scenario 1), in which a range expansion occurs into a previously empty landscape, we implemented scenarios that include linearly increasing spatial gradients in local mortality (scenarios 2 and 3). We contrast a setting in which dispersal propensity may evolve and the organisms do not have the capacity to sense the environmental change in such a gradient (scenario 2) and a scenario in which we assume that individuals have perfect information to make an optimal dispersal decision plastically and therefore evolutionary changes become irrelevant (scenario 3). Informed dispersal is based on a cost-benefit analysis, which takes into account population densities (i.e., competition) in the patch of origin and in all potential target patches, as well as the effect of the mortality gradient.

The model was designed to be as simple as possible and to provide qualitative predictions on the impact of environmental gradients and information use on the ecological and evolutionary dynamics of range expansions. We therefore ran an extensive sensitivity analysis (Figs. S6 – S10). We neither parametrize nor fit the model to the experimental data.

#### Landscape and the environmental gradient

For simplicity, we assume a linear landscape of 100 interconnected patches. At the start of each replicate simulation only the first five patches are populated. The landscape allows individuals to disperse following a stepping stone model, that is, we assume nearest neighbor dispersal with reflecting boundary conditions at both ends of the landscape. In scenarios 2 and 3, which include an environmental mortality gradient, we assume that this additional source of local mortality (*μ_x_*) acts after reproduction and density regulation (see below) and before dispersal. The mortality gradient is linear and increases from *μ*_1_ = 0 in the first patch to *μ*_100_ = 1 mortality in the last patch.

#### Dispersal

Besides being governed by the landscape setting as described above, dispersal of individuals is assumed to be either genetically controlled (scenarios 1 and 2) or fully plastic and informed (scenario 3). We here only describe the former two scenarios, the latter will be dealt with in detail below. The probability of dispersing, more specifically emigrating from a natal patch, is genetically controlled by a haploid locus that codes for the dispersal rate (*d_i_*). When an individual (*i*) disperses according to its specific dispersal rate, the direction in the linear landscape (i.e., towards the range core or towards the range front) is drawn randomly.

We do not assume explicit dispersal costs (Bonte *et al.*, 2012). However, dispersal is implicitly costly, as we assume that dispersal trades off with reproduction and competitive ability as described below.

#### Reproduction and density regulation

Reproduction occurs after dispersal and follows a modified logistic, density-dependent growth model based on Beverton & Holt (1957):

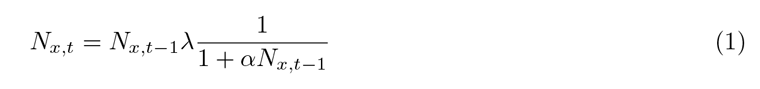

where *N_x,t_* is the population size in patch *x* at time *t*, λ is the growth rate and *α* the intra-specific competition coefficient as introduced above. As reproductive (λ*_i_*) and competitive ability (*α*_i_) are individual-based traits, the mean number of offspring an individual produces at time *t* in a population of size *N_x,t_* is:

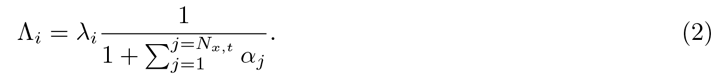

We include demographic stochasticity by assuming that reproduction follows a Poisson process and drawing the realized number of offspring for individual *i* from a Poisson distribution with mean Λ*_i_*. After reproduction all individuals of the previous generation die.

#### Trait correlations and trade-offs

As outlined in the Electronic Supplementary Material (“Linking consumer-resource dynamics to logistic growth”), we assume that reproductive and competitive ability (λ*_i_* and *α_i_*, respectively) are individual-based traits that correlate positively:

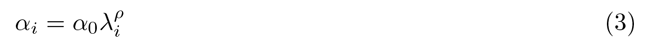

with *α*_0_ as a baseline competitive ability and *ρ* as the correlation exponent between competitive and reproductive ability. As Fronhofer & Altermatt (2015) showed previously, a large part of changes in competitive ability seem driven by changing feeding rates and not by changing assimilation coefficients. We therefore assume *ρ* = 2 as a standard scenario following the logic outlined above. For a summary of parameters and tested values refer to Tab. S1.

Furthermore, we assume that dispersal is costly (Bonte *et al.*, 2012) and trades off with reproduction, and, therefore, also with competitive ability:

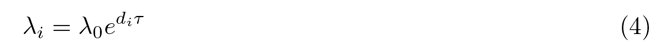

where λ_0_ is the baseline fecundity, *d_i_*, the dispersal rate of individual *i* and *τ* the strength of the trade-off between dispersal and fecundity.

#### Information use

In scenario 3 we assume that dispersal is plastic in the sense that individuals make informed dispersal decisions. The decision of whether to disperse to one of the two neighboring patches in the linear landscape or to stay in the natal patch is based on a cost-benefit calculation. We assume that individuals have perfect knowledge on the patch densities in their natal patch (*N_x_,_t_*) and in the potential target patches, as well as information on local mortality (*μ_x_*) due to the mortality gradient. Individuals disperse to the patch x that maximizes their fitness according to Eq. 2:

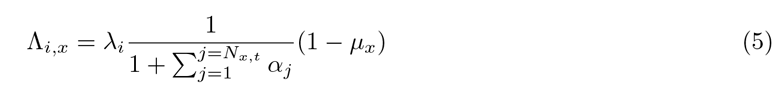

This approach only accounts for direct fitness benefits and ignores inclusive fitness (Hamilton & May, 1977). Our simulations therefore underestimate dispersal and spatial spread rates in the informed scenario. For a detailed treatment of the effect of kin competition on range dynamics see Kubisch *et al.* (2013).

#### Evolution and the genetic algorithm

Evolutionary dynamics are an emergent phenomenon of any individual-based model that allows for variation in heritable, individual-based traits. The specific simulation scenario leads to selection pressures, such as spatial selection (Phillips *et al.*, 2010; Shine *et al.*, 2011) in range expansion scenarios, for instance. We here assume that dispersal rate (*d_i_*), fecundity (λ*_i_*) and competitive ability (*α_i_*) are heritable and passed on from parent to offspring with a mutation rate *m* = 0.001 that leads to a random change of the trait value drawn from a Gaussian distribution with mean zero and standard deviation Δ*m* = 0.1. The only trait subject to mutations is the dispersal trait (*d_i_*) since both fecundity (λ*_i_*) and competitive ability (*α_i_*) depend on dispersal via the trade-off and correlation structures explained above (Eq. 3 and 4).

At the genotype level we do not implement any boundary conditions on the dispersal trait, that is, depending on mutations *d_i_* may be negative or > 1. At the phenotype level values < 1 are set to zero and values > 1 are set to 1. These phenotypic values are are also used to calculate fecundity according to Eq. 4.

#### Simulation experiments

All simulations were initialized with populations at a baseline equilibrium density 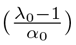 in the first five patches in order to allow the individuals to subsequently spread through the landscape. Individuals in these populations were initialized with random dispersal rates (0 ≤ *d_i_* ≤ 1) as standing genetic variation. All simulations were allowed to proceed for 95 generations which, given the stepping stone dispersal model, is the minimal time span needed to reach the opposite end of the landscape. In general, simulations were replicated 20 times. The sensitivity analysis of scenario 3 (gradient and information) was performed on less replicates (between 1 and 10) as these simulations show only very little variation between replicates (see Fig. 1 E) and take an excessive amount of time to run. Please see Tab. S1 for tested parameter combinations and Figs. S6 – S10 for a sensitivity analysis.

### Microcosm Experiments

#### Study organism

We used *Tetrahymena pyriformis,* a unicellular freshwater ciliate, as a model organism (Altermatt *et al.*, 2015; Fronhofer & Altermatt, 2015). *Tetrahymena pyriformis* is small (approx. 40-50 ym along the major axis), has a relatively short doubling time (approx. 4-5h) and reaches high densities (equilibrium densities: 5,000-15,000 individuals/mL) which makes it well suited for ecological and evolutionary experiments (Altermatt *et al.*, 2015). We kept *T. pyriformis* under controlled environmental conditions at 20ΰC in protist pellet medium (0.46 g/L; Carolina Biological Supply) with bacteria (5 vol-% of standardized 7-day-old cultures of *Serratia fonticola, Brevibacillus brevis* and *Bacillus subtilis*) as food resources. We used the same protist cultures as Fronhofer & Altermatt (2015) and therefore started evolution experiments with standing genetic variation. The cultures were originally obtained from Carolina Biological Supply and regularly restocked to conserve genetic variation (Cadotte, 2007).

#### Microcosm landscapes

The range expansion experiments were performed in linear landscapes consisting of 14 interconnected microcosms (patches). We used 20 mL vials (Sarstedt), connected them with silicone tubing (VWR; 4mm inside diameter) and a stopcock (B. Braun Discofix) to regulate dispersal (length of tubing and stopcock: 6 cm). All experiments were replicated 6 times in two experimental blocks of 3 replicates each separated by 1 day.

#### Scenarios and experimental procedure

At the beginning of each experiment, the first patch of a landscape was filled with a week-old *Tetrahymena pyriformis* culture that had reached its equilibrium density. Subsequently, the stopcocks were opened and dispersal was allowed for 4 hours. In order to avoid aging of medium and to limit contaminations, the landscape was not completely filled with medium from the start of the experiment, but empty patches were added subsequently to the landscape front. At the beginning of the experiment, 3 of the 14 patches were filled. At each day of the experiment, one additional patch filled with freshly bacterized medium (5 vol-%) was added at the front. Since all patches between range core and range front were connected, dispersal could potentially occur across multiple patches and towards the range front as well as towards the range core.

To analyze the influence of information use on the eco-evolutionary dynamics of range expansions into environmental gradients, we designed two experimental treatments in addition to the control treatment (scenario 1) described above. For both, uninformed (scenario 2) and informed scenarios (scenario 3) a linear mortality gradient was applied, ranging from 0% mortality in the first patch to 100% mortality in the last patch. In the uninformed scenario (scenario 2), depending on the mortality gradient, a certain volume of the microcosm was removed, discarded and replaced with bacterized medium. In the informed scenario (scenario 3), we followed the same procedure but replaced the volume with dead *T. pyriformis* from a 4-days old culture that was killed by ultrasonication (duration: 4 min.; amplitude: 40%; pulse on: 2 sec.; pulse off: 1 sec; ice bath to avoid heating). We therefore use dead *T. pyriformis* and their chemical cues to inform the protists in the experiments about the increasing mortality in the landscape. Previous to the experimental evolution assays we performed chemical orientation assays to confirm that dead conspecifics are indeed used as a negative tactic cue (see Electronic Supplementary Material “Effects of chemical cues provided by dead conspecifics”).

The general experimental procedure was as follows: we first applied the respective treatments (scenario 1: control, scenario 2: mortality gradient, scenario 3: mortality gradient and information) and allowed for dispersal (4h) on one day. The following day allowed for regrowth. We therefore had discrete dispersal and growth phases in analogy to the individual-based model described above. In total, the evolution experiment took 26 days with 13 dispersal events and subsequently two days of common garden. Each scenario was replicated 6 times and the experimental units were arranged in two blocks of 3 replicates each shifted by one day due to the large number of samples to process.

#### Common garden and growth curves

In order to tease apart plastic changes, due to environmental or parental effects, in dispersal (respectively, movement strategies), growth rates and competitive abilities from genetically or non-genetically inherited evolutionary changes, we transferred range core and range front populations to a common environment after the experimental evolution phase. We transferred all core and front populations from the end of the experiment to 200 mL Erlenmeyer flasks and added 100 mL freshly bacterized medium to the 15 mL from the experimental microcosms. This transfer reset all populations to roughly the same environmental conditions in terms of resource availability and chemical composition of the medium. After 2 days in this common environment, all populations were assessed for divergence in movement behaviour, growth rates and competitive abilities.

Growth rates and competitive abilities were estimated by performing growth curve experiments and subsequently fitting logistic growth curves (Eq. S2) to the time-series data. All growth curves were started with approx. 500 individuals per mL by diluting the populations from the common garden. 5 vol-% bacteria from a standardized, 7-days old culture were added as resources. The growth of each population was followed for 10 days using video recording and analysis as described below.

Logistic growth curves were fitted to the individual replicates using a least-squares approach. Eq. S2 was solved (function ‘ode’ of the ‘deSolve’ package in R version 3.2.3) and the model was fit using the Levenberg-Marquardt algorithm (function ‘nls.lm’ of the ‘;minpack.lm’ package) which minimizes the sum of squared residuals.

#### Data collection

Before a treatment was performed, a 0.5 mL sample of each patch was collected. In the control and uninformed scenario, the sampling volume was replaced with fresh, bacterized medium. In the informed scenario, the sampling volume was replaced with dead *T. pyriformis* and fresh, bacterized medium for the first patch, respectively.

A subsample was then used for video recording with a Leica M205 C stereomicroscope (16 fold magnification) and a Hamamatsu Orca Flash 4 video camera (imaged volume: 34.4 *μ*L; sample height: 0.5 mm). Videos of 20 seconds were recorded with a total of 500 gray scale images with a resolution of 2024 × 2024 pixels.

The general method of automated image analysis was introduced by Pennekamp & Schtickzelle (2013); Pennekamp *et al.* (2015) and has successfully used in previous experiments (Giometto *et al.*, 2014; Fronhofer *et al.*, 2015a; Fronhofer & Altermatt, 2015; Fronhofer *et al.*, 2015b). The aim is to collect abundance data as well as morphological and behavioral data simultaneously and provide information at the individual level. The principle of automated image analysis first includes a cleaning step followed by different analytical steps to determine morphological traits (length, size), abundance and movement data (velocity, turning angle, Euclidean distance). The first step of the image analysis consists in identifying the objects of interest by segmenting the moving foreground from the static background. Therefore the difference between picture t and t + 1 was analyzed. In general, only particles with a size between 20 and 200 pixels and a minimal path length of 100 frames were included in the analysis. Trajectories of each individual were analyzed with the ImageJ MOSAIC plugin (Sbalzarini & Koumoutsakos, 2005). Data of each sample (abundance, velocity, body size, turning angle) was saved as mean values. As previous work consistently showed that dispersal rates and movement behaviour correlate highly in these protist microcosms (Fronhofer & Altermatt, 2015; Fronhofer *et al.*, 2015b), we here use movement as a proxy for dispersal. Data can be downloaded from Dryad DOI: XXX.

#### Statistical analysis

Differences in velocity were analysed using linear mixed models (LMM). We included the experimental block (replicates 1-3 and 4-6) as a random effect in our analyses. We used a Gaussian error structure as the QQ-plots indicated that this assumption was not heavily violated. All analyses were performed at the population level, i.e. on mean parameters over all individuals in a sample. This approach is very conservative, since it significantly reduces the sample size given the high population densities and the individual-based data collected by video recording and analysis. These analyses were performed using R version 3.2.3 and the “lmerTest” package.

The distribution of population densities over space was compared between treatments using the empirical cumulative population density distributions (see Fig. S3 D–F). Again, we chose a very conservative approach and only compared the median cumulative density distributions of the treatments using the Cramer-von Mises (CvM) statistic (*ω*^2^) for two samples. We therefore calculated the sum of the squared differences between two empirical cumulative density distributions (*ω*^2^). We subsequently analysed significance levels by resampling (one-sided tests) and additionally provide Probability-Probability plots for visual analysis (Fig. S4). As we performed all pairwise comparisons (2 comparisons per treatment), we corrected the obtained significance thresholds using the Bonferroni method, which consists of multiplying the initially obtained significance thresholds with the number of comparisons.

The chemical orientation assay was analysed using generalized linear mixed models (GLMM) with binomial error distributions and counts of individuals choosing either the treatment or the control patch. We included “replicate” as a random effect to take into account the pairing between dispersal to control and treatment patches within one replicate. We further included a sample level random effect to account for overdispersion.

The empirical correlation between competition coefficients (α) and growth rates (*r*_0_) for populations from the range core and the range margin was analysed using non-linear regressions (following Eq. 3) for grouped data with the function “nlsList” of the “nlme” package in R version 3.2.3. For this analysis, we only used data from scenarios 1 and 2 as we did not observe evolutionary dynamics in scenario 3 so that the classification into core and front populations is not meaningful. We nevertheless report data from scenario 3 in Fig. 3 B.

## Results

### Theoretical predictions

In the control and gradient scenarios our theoretical analyses (Fig. 1) predict evolutionarily increased dispersal at the range front compared the range core (Fig. 1 B, D). However, the difference in evolved dispersal propensities between range core and front populations is reduced in the gradient scenario. Furthermore, we predict higher population densities at range fronts in the control scenario and, to a lesser extent, also in the gradient scenario (Fig. 1 A, C and S3 A, B). The invasion does not proceed as far in the gradient scenario as in the control, suggesting that a stable range border forms (Fig. 1 A, C and S3 A, B).

In the informed dispersal scenario, the density profile of populations across the range is inverted in comparison to the evolutionary scenarios implying lower densities at range fronts in comparison to range cores (Fig. 1 E and S3 C). These predictions qualitatively hold true across a large range of tested parameter values (Tab. S1; Figs. S6 – S10; especially for weak dispersal-fecundity trade-offs and fecundity-competition correlation coefficients > 1).

### Experimental range dynamics

Our experimental results corroborate our theoretical predictions (Fig. 2). At the end of the range expansion phase we found increased movement velocities (which correlate strongly with dispersal (Fronhofer & Altermatt, 2015; Fronhofer *et al.*, 2015b)) at range fronts (Fig. 2 B, E, H), although the effect was weak in the informed scenario (control: LMM, space: *N* = 74(6), *df* = 72 *t* = 11.79, *p* < 0.001; gradient: LMM, space: *N* = 77(6), *df* = 74 *t* = 13.24, *p* < 0.001; information & gradient: LMM, space: *N* = 64(6), *df* = 62 *t* = 4.69, *p* < 0.001). After the common garden, the velocities in range core, respectively range front populations, were still significantly different in the control (Fig. 2 C; LMM, range position: *N* = 12, *df* = 9 *t* = 3.94, *p* = 0.0034) and in the gradient scenario (Fig. 2 F; LMM, range position: *N* = 12, *df* =10 *t* = 7.23, *p* < 0.001). No differences were observed in the informed scenario (Fig. 2 I; LMM, range position: *N* =12, *df* =10 *t* = −0.045, *p* = 0.965).

Furthermore, we observed the predicted spatial distribution of population densities with high densities at range fronts and low densities in range cores in the control and gradient scenario (Fig. 2 A, C and S3 D, E). Information use completely inverted this pattern leading to significantly different distributions of population densities between informed and uninformed scenarios (Fig. 2, S3 D–F and Fig. S4)

**Figure 1:**
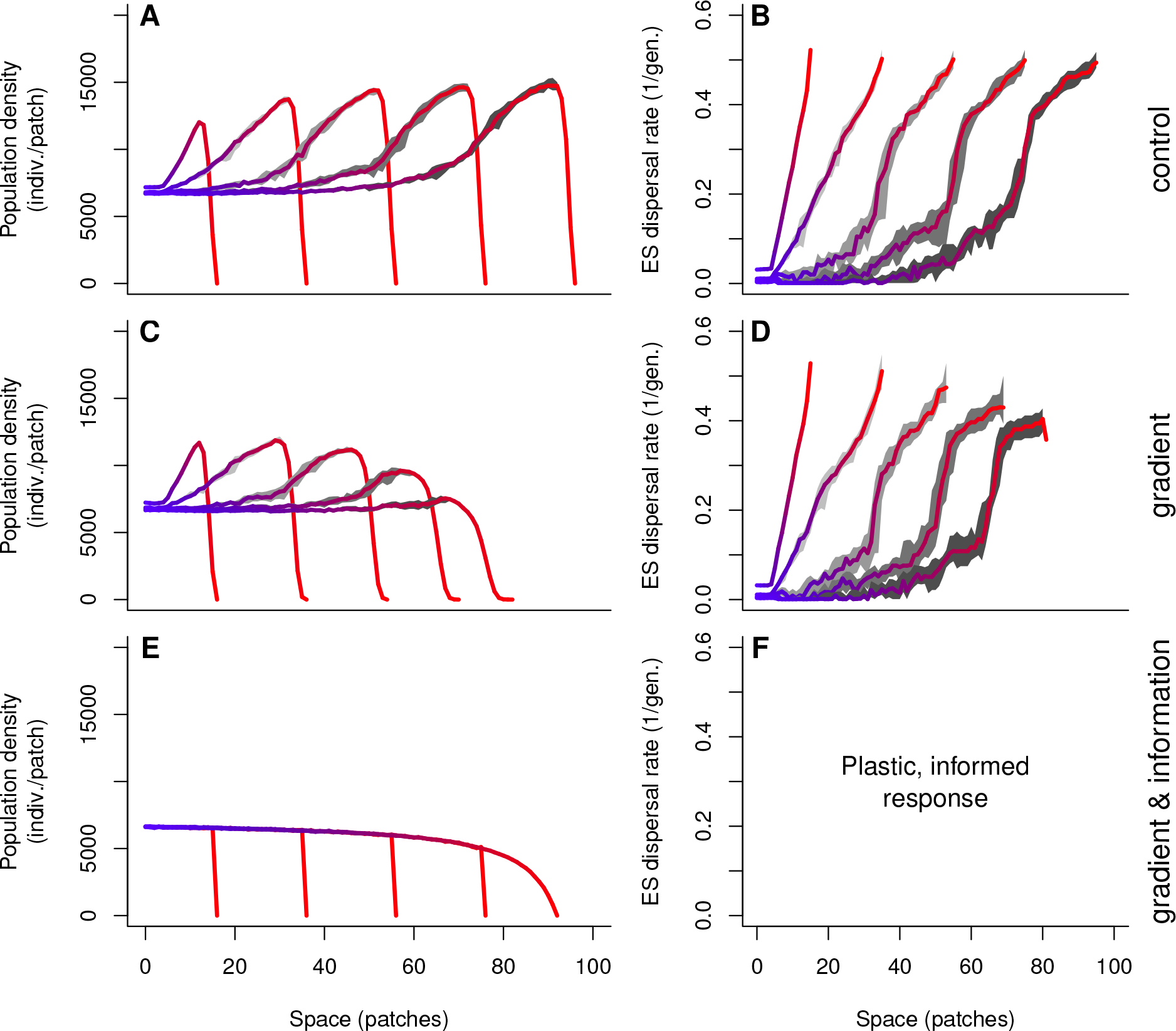
Range dynamics — theoretical predictions. (A-B) Expansion into homogeneous environment (control). Population densities increase from range core to front due to dispersal-fecundity trade-offs (Eq. 4) and fecundity-competition correlations (Eq. 3). Spatial selection leads to increased dispersal at range fronts. (C-D) Expansion into a mortality gradient. Density patterns are not fundamentally altered during a major part of the expansion (see also Fig. S3). However, increasing mortality locally reduces population densities and selects against dispersal. (E-F) Expansion in a mortality gradient and information use. Dispersal is plastic and individuals are fully informed about the mortality gradient, population densities in their natal and potential target patches (Eq. 5). The distribution of population densities over space is inverted (Fig. S3). Dispersal does not evolve, but it is predicted to be plastically higher at the range front during the expansion due to the decision rule. Temporal snapshots: *t* = [10, 30, 50, 70, 90]. Parameter settings: λ_0_ = 14, *α*_0_ = 0.00001, *ρ* = 2, *τ* = 2. We report medians over 20 replicate simulations (solid line; blue (range core) to red (range front)) and the 25th and 75th percentiles (grey shading; darker with time).

### Concurrent changes in reproduction and competition

At the end of the experiment, we measured population growth rates and competitive abilities after a common garden phase to separate genetic from plastic effects. We observed a positive correlation between growth rate and competitive ability (Fig. 3 B), corroborating our assumption about this correlation (Fig. 3 A; for details see Eq. 3 and the Electronic Supplementary Material). While individuals from range cores followed the theoretically predicted correlation quantitatively, individuals from range fronts shifted the predicted correlation curve towards increased growth rates (Fig. 3 B).

**Figure 2:**
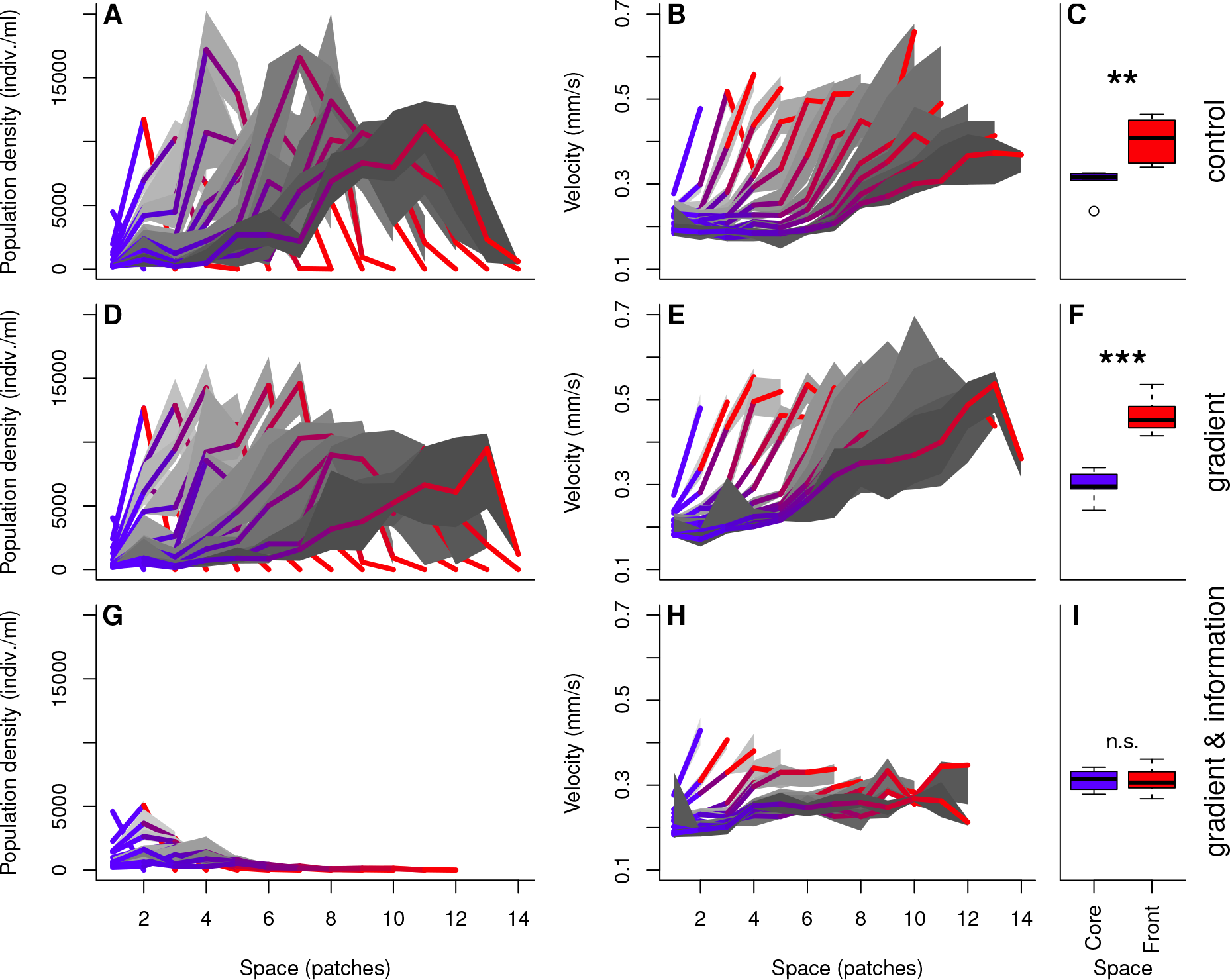
Range dynamics — experimental results. As predicted, the spatial distribution of population densities (A, D, G) showed an increase in densities towards the range front in the control (A) and gradient (D) scenarios. Information use (G) inverted this pattern (see Fig. S3 D–F; the distribution is statistically different from the other two; Fig. S4). On the last day of the evolution phase clear differences in movement over space was found in all scenarios (B, E, H), although the effect was weak in the informed scenario. After the common garden phase, the velocities in range core respectively range front populations were still significantly different in the control (C) and in the gradient scenario (F). No differences were observed in the informed scenario (I). We report medians over 6 experimental replicates (solid line; blue (range core) to red (range front)) and the 25th and 75th percentiles (grey shading; darker with time). Stars indicate statistical significance (see text for details).

**Figure 3:**
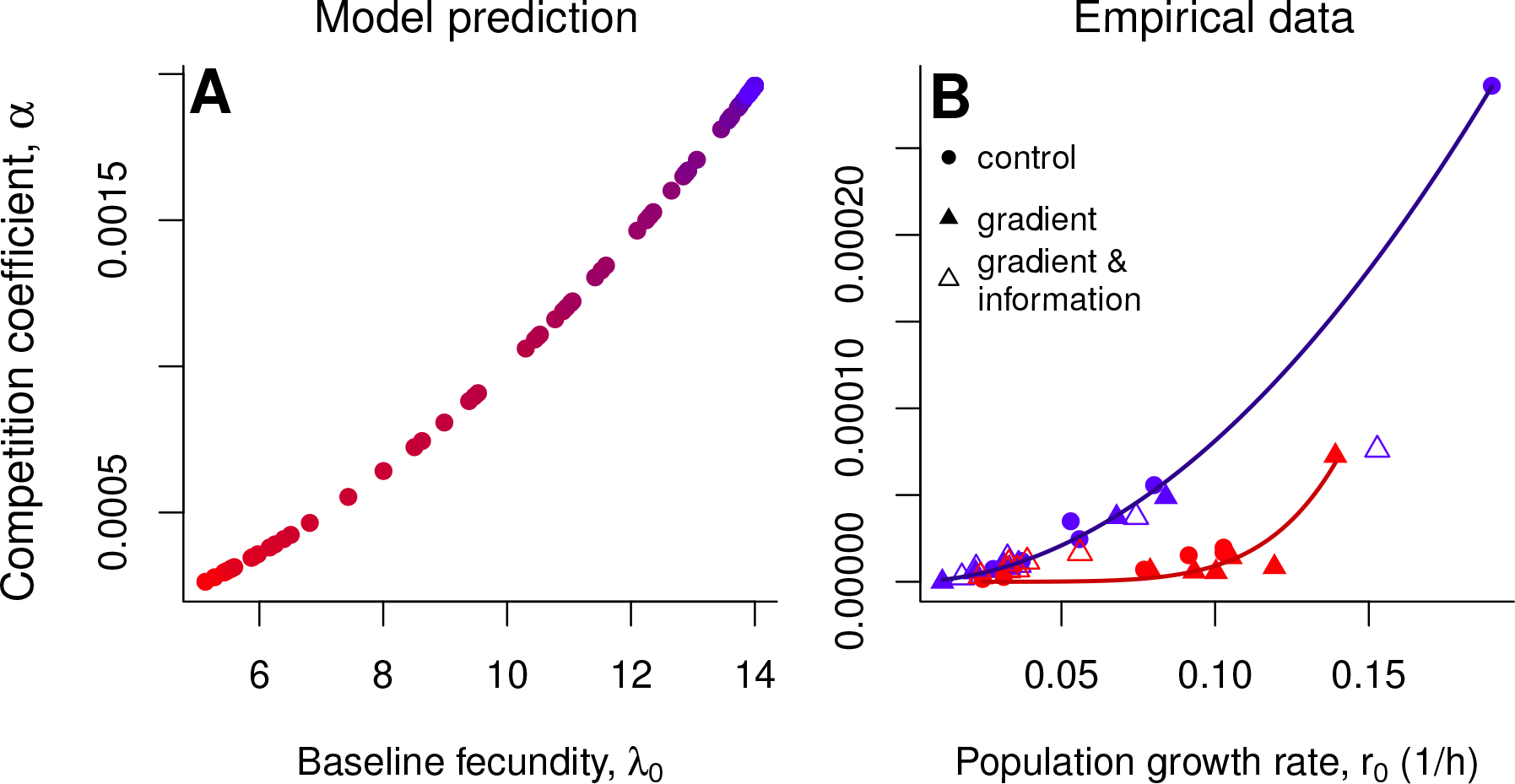
Concurrent evolution of reproduction and competition. (A) As derived in the Electronic Supplementary Material, our model assumes a correlation between competitive ability (*α*) and fecundity (λ; Eq. 3). Given a linear functional response we predict a roughly quadratic relationship (λ–*α* correlation coefficient *ρ* = 2). Due to the trade-off between dispersal and fecundity, high fecundities and competitive abilities are predicted in the range core, where individuals are less dispersive (blue color tones indicate range core and red color tones range front; data from the control scenario; see Fig. 1). (B) Empirically measured competition coefficients (*α*) and growth rates (*r*_0_) after the common garden phase. The theoretically predicted relationship between competition and reproduction was found for core populations (blue; empirically measured *α*_0_ = 0.0074 (CI: 0.0053, 0.0094); *ρ* = 1.96 (CI: 1.80, 2.11); only data form scenarios 1 and 2). However, selection acting during the range expansion altered this relationship (red; *α*_0_ = 11.24 (CI:-29.11, 51.60); *ρ* = 6.09 (CI: 4.31, 7.86); only data form scenarios 1 and 2) allowing individuals at the range front to have higher reproductive rates than theoretically predicted. Increased reproduction is highly advantageous as populations at the range front experience strong selection for both dispersal and reproductive ability.

## Discussion

Hitherto, research on range dynamics has often assumed homogeneous environments and consistently ignored that universally occurring environmental gradients provide information to spreading organisms about local conditions. This information may allow spreading organisms to plastically adapt their dispersal decisions which has the potential to alter macroecological patterns. We now theoretically and experimentally show that the ecological and evolutionary dynamics of species’; ranges are not only driven by the direct, fitness relevant effect of environmental gradients but, most importantly, by the information content of such gradients.

We find that range expansions lead to increased dispersal at the range front in the control and gradient scenarios (Fig. 2 C, F), which is consistent with previous theoretical (Kubisch *et al.*, 2014), comparative (Phillips *et al.*, 2006) and experimental results (Fronhofer & Altermatt, 2015). Importantly, however, the latter has hitherto only been studied in unrealistic environmentally homogeneous landscapes. The evolutionary increase in dispersal is due to spatial assortment and fitness advantages of dispersers that colonize empty habitat at the range front and therefore do not suffer from competition (“spatial selection” Phillips *et al.* 2010; Shine *et al.* 2011). In the gradient scenario, spatial selection is counteracted by increasing mortality. In the informed scenario, we find differences in dispersal only early during the range expansion phase, but not after the common garden (Fig. 2 H, I), which confirms our model assumption regarding complete plasticity of dispersal in this scenario.

Interestingly, theoretical predictions and experimental results show a spatial density pattern of increasing population sizes towards the range front (Figs. 1, 2). Thus, information use inverts the spatial distribution of population densities across a species’; range. These density patterns emerge in the theoretical results because more dispersive individuals at range fronts deplete resources less due to the trade-off between dispersal and reproduction (Eq. 4) and concurrent changes in competitive abilities (Eq. 3), which implies that patches at the range front can support higher equilibrium population densities (Fronhofer & Altermatt, 2015). Our empirical results, especially the observed correlation between reproduction and competition (Fig. 3), support our model assumptions and the relationship between growth rate and competitive ability derived in the Electronic Supplementary Material.

Remarkably, in our experiments, the impact of the mortality gradient on the spatial distribution of densities was relatively weak, while the influence of information use was extremely strong in inverting the spatial pattern of population densities (Fig. 2 D, G and S3 D–F). This indicates that not the environmental gradient itself, but rather using information thereon drives range expansion dynamics into environmental gradients.

In our experiments, the mortality gradient selected for increased reproduction (Fig. 3 B). The quantitative difference between theoretical prediction and experimental results (Fig. S3) regarding the impact of information use can be linked to the shift in trait correlation structure we observed (Fig. 3). This shift in the correlation can be interpreted as the result of strong selection for high reproduction at range fronts, which explains why the effect of the mortality gradient was relatively small in the experiments (Fig. 2 D): Populations overcame increased mortality by increasing reproduction. The shift in the trait correlation structure is likely due to a change in foraging behavior from a linear to a saturating functional response (see Electronic Supplementary Material) as reported previously (Fronhofer & Altermatt, 2015).

In conclusion, we show that environmental gradients indeed have a two-fold effect consisting of 1) a direct fitness relevant effect of the gradients itself and 2) of the information the gradient conveys on the environmental change. This information can be used to inform dispersal decisions which has major consequences for the macroecological patterns of range expansions along environmental gradients. Informed dispersal does not only impact expansion dynamics but can completely invert the spatial distribution of population densities. Our theoretical and experimental findings highlight the need to include environmental heterogeneity and organisms’ capacity to process information thereon into realistic predictions of invasion dynamics and range expansions.

## Acknowledgements

We thank Mark van Kleunen for his input during the planning of experiments, Sebastian Schreiber for his literature suggestions and Anita Narwani for feedback on an earlier version of the manuscript. Funding is from Eawag (to E.A.F.) and the Swiss National Science Foundation, Grant No. PP00P3_150698 (to F.A.).

## Author contributions

E.A.F., N.N. and F.A. designed the research; N.N. performed the experiments; N.N. and E.A.F. analysed the data; E.A.F. developed the stochastic modelling framework; E.A.F. and F.A. wrote the paper.

## Data accessibility

All data and computer code will be archived in Dryad and the DOI will be included at the end of the manuscript.

## Electronic Supplementary Material

### Linking consumer-resource dynamics to logistic growth

Although logistic growth models are not mechanistic in the sense that they are abstract and descriptive representations of population dynamics, such simple models may be useful and tractable approximations of ecological dynamics. However, the use of such models, especially of the classically used *r* — *K* formulation,

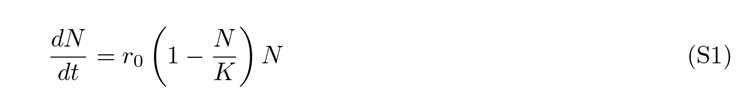

may lead to erroneous predictions since, due to the level of abstraction, the parameters of the model are difficult to interpret in biologically relevant terms. This is especially true for “carrying capacity” (*K*) and ideas linked to “*K*-selection” (Matessi & Gatto, 1984; Mallet, 2012; Reznick et al., 2002; Rueffler et al., 2006). Consumer-resource models are an alternative and more mechanistic framework, which does not have the same limitations and the biological interpretations are more direct. Specifically, they can be used in an eco-evolutionary context since model parameters linked to resource use (e.g., search efficiency, handling time) are related to real traits that can be subject to evolutionary change (Fronhofer & Altermatt, 2015).

The well known downside of more mechanistic models is their often increased complexity, especially in terms of number of parameters (for instance, moving from 2 parameters for the logistic to 5 to 7 depending on the consumer-resource model). In the following, we will use previous work by MacArthur (1970), Schoener (1973) and Abrams (2009) to lay out how one can use a consumer-resource model to derive a logistic model of population growth (Matessi & Gatto, 1984), which, in contrast to the *r* — *K* formulation (Eq. S1), has more biologically relevant and interpretable parameters (Mallet, 2012). Importantly, this approach enables us to generate specific hypotheses regarding how natural selection, acting on individual traits linked to foraging activities (search efficiency, handling time), can alter more abstract, population level model parameters.

A central issue with the logistic model introduced above (Eq. S1) is linked to the “carrying capacity” parameter (*K*; for a detailed discussion see Mallet 2012). Therefore, we will here use the original, *r* — *α* formulation of the logistic equation proposed by Verhulst (1838):

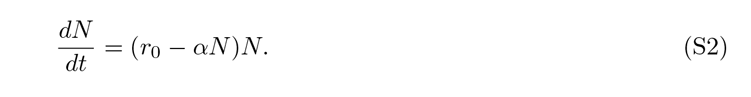

In this formulation *r*_0_ is the intrinsic rate of increase and *α* represents the intraspecific competition coefficient. The equilibrium density is given by 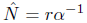. In the logistic model provided by Eq. S2 the per capita growth rate of the population is

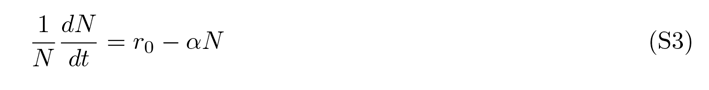

implying that density dependence acts in a linear way with *r*_0_ being the maximal growth rate when population size (*N*) is small. The growth rate then decreases as a function of *N* with slope α.

As a consumer-resource model, we use a Lotka-Volterra model with a linear functional response (Holling Type I) and logistic resource growth. The consumer dynamics (*N*) are described by:

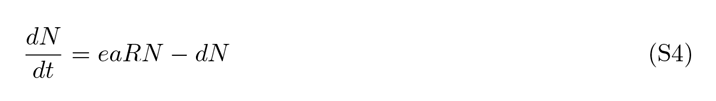

with *e* as the efficiency of converting resources into consumers, *a* as the feeding rate, that is, the slope of the linear functional response, *d* as the consumer’s mortality rate and *R* as the resource population size. The per capita formulation then is:

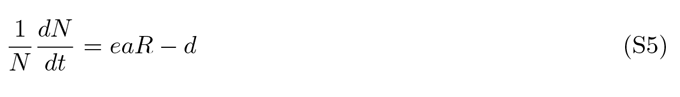

The resource dynamics (R) follow:

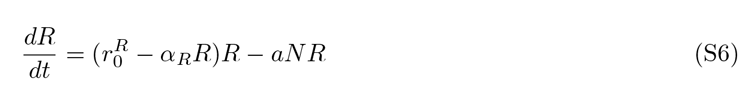

with 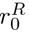 as the resource’s intrinsic rate of increase and *α_R_* as its competition coefficient in analogy to Eq. S2.

We can now reformulate Eq. S3 slightly to make the parallels to Eq. S5 more obvious using the fact that *r*_0_ = *b*_0_ – *d*_0_ with *b*_0_ as birth and *d*_0_ as death rate which yields 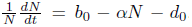. As a consequence of equating this formulation with Eq. S5 we can write

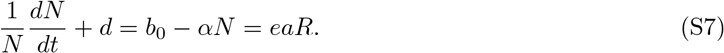

Knowing *R* allows us to understand how the consumer’s growth rate (*r*_0_) and competition coefficient (*α*) are related to its more mechanistically based traits, including efficiency (*e*) and feeding rate (*a*). This is classically done by assuming that the resources are always at equilibrium, that is, 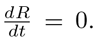. From Eq. S6 we obtain that either *R* = 0, which is the trivial case and not under further consideration, and
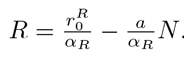. Including this information into Eq. S7 yields:

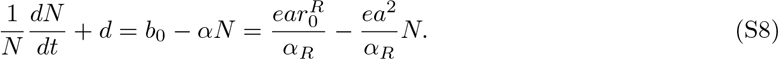

Note that the relationship is not fundamentally changed by a Type II functional response. The overall slope would still be negative, however the shape of density-dependence becomes non-linear as in the *θ*-logistic model.

This relationship allows us to reach the following conclusions: keeping resource dynamics constant, the consumer’s birth rate (*b_0_*) and, by extension (if the death rate is sufficiently small), its intrinsic rate of increase (*r*_0_), linearly increases with both the consumer’s efficiency (*e*) and its feeding rate (*a*). The competition coefficient (*α*) increases linearly with efficiency (*e*) and quadratically with feeding rate (*a*). Consequently, the intrinsic rate of increase (*r*_0_) and the competition coefficient (*α*) should correlate positively.

From an evolutionary point of view, Matessi & Gatto (1984) show that density-dependent selection (“*K*-selection”) should, rather than maximizing the equilibrium density, minimize equilibrium resource availability and consequently 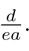. Therefore, density-dependent selection, as individuals might experience in range core populations, can be predicted to increase *r*_0_ but especially *α* very strongly, as the relationship with feeding rate is quadratic for the latter parameter. Individuals in those same populations are expected to show reduced dispersal rates if the habitat is stable. Expanding this logic into possible trade-offs with dispersal ability, we can justify that, as hypothesized by Fronhofer & Altermatt (2015), more dispersive individuals, as one can find at range fronts for instance, should have lower feedings rates and efficiencies. Consequently, we can assume to find a (negative) trade-off between dispersiveness and *r*_0_, respectively *α*.

### Effects of chemical cues provided by dead conspecifics

#### Chemical orientation and movement

To investigate whether dead individuals of *T. pyriformis* were an appropriate source of information for their living conspecifics, a chemical orientation assay with different concentrations of dead individuals was performed prior to the evolution experiments. A three-patch system was used, consisting of three 20 mL vials (Sarstedt) connected by silicon tubing (VWR, 4 mm inner diameter) and a three-way stopcock (Braun, Discofix). The patch of origin was filled with 15 mL of a 4-day-old *T. pyriformis* culture. One of the target patches was filled with bacterized medium, the other target patch was filled with 10 %, 50 % or 100 % dead *T. pyriformis,* respectively. To obtain dead *T. pyriformis,* a culture of 100 mL was killed by ultrasonication for 4 minutes (amplitude: 40 %, pulse on: 2 sec, pulse off: 1 sec).

Dispersal form the patch of origin to either target patches was allowed for 4 hours. To assess the density of *T. pyriformis* after dispersal, all patches were analyzed using video recording and analysis as described below. The experiment was replicated 10 times for each concentration of dead *T. pyriformis.* The results of the chemical orientation assay are shown in Fig. S1 and confirmed that chemical cues from dead conspecifics are used as a cue for negative taxis.

#### Effects on population growth

Besides being used as a source of information, the chemical compounds from dead conspecifics may also have direct negative effects on *T. pyriformis* movement behavior, growth and competitive abilities. In order to take these potential side effects into account, we quantified the impact of different concentrations of dead conspecifics by recording growth curves, i.e. time series of growing *T. pyriformis* populations initially exposed to 0, 10, 50 and 75 vol-% of sonicated, dead conspecifics form a 4-days old culture that had reached its equilibrium density. We subsequently fit logistic growth curves to the time series as described in the main text. The results are reported in Fig. S2 and indicate that growth is impacted negatively by chemical cues from dead conspecifics.

In order to explore potential consequences of this negative effect on range dynamics, we ran additional simulations which explored theoretically the dynamics of range expansions into combined mortality and decreasing fecundity gradients (Fig. S5). These results indicate that, while range expansions may proceed more slowly, the consequences of decreased fertility and information use are qualitatively different: only information use is predicted to invert the spatial cumulative population density distribution as reported in Fig. S3 A–C. Note that reduced growth rates may also be due to increased mortality. The latter case would increase the steepness of the mortality gradient but not change results qualitatively.

## Supplementary figures

**Supplementary Figure S1:**
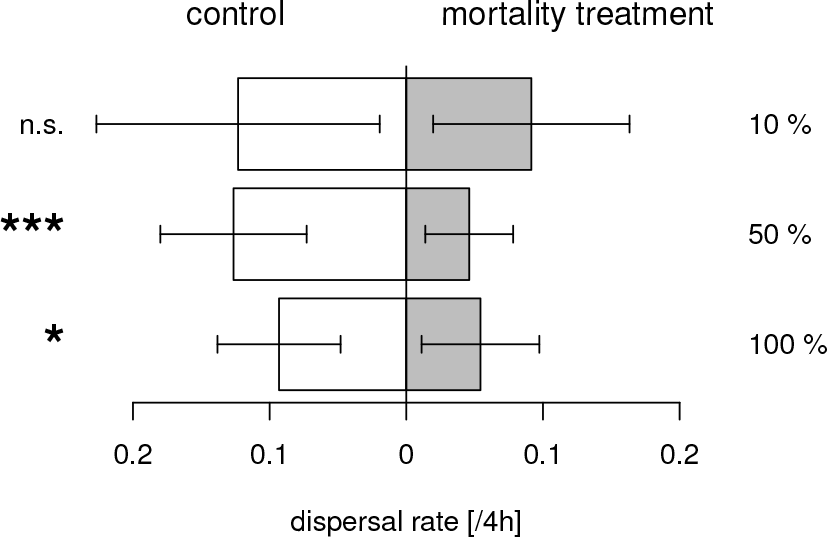
*Tetrahymena pyriformis* — chemical cues of dead conspecifics used for negative chemotaxis. Choice experiments indicate that *Tetrahymena pyriformis* indeed avoid patches with dead conspecifics if the concentration of chemical cues is high enough, i.e., over 10 % (concentrations tested: 10%, 50% and 100%). The bars (mean *±* s.d.; 10 replicates) indicate the dispersal rate from the patch of origin to either of two target patches (control with fresh medium vs. mortality treatment with varying concentrations of dead conspecifics). Stars indicate significance levels: GLMM (treatment = 10%): *N* = 12263(10), *z* = —0.81, *p* = 0.42, error family: binomial; GLMM (treatment = 50%): *N* = 9798(10), *z* = −3.63, *p* < 0.001, error family: binomial; GLMM (treatment = 100%): *N* = 6546(10), *z* = −2.31, *p* = 0.021,error family: binomial.

**Supplementary Figure S2:**
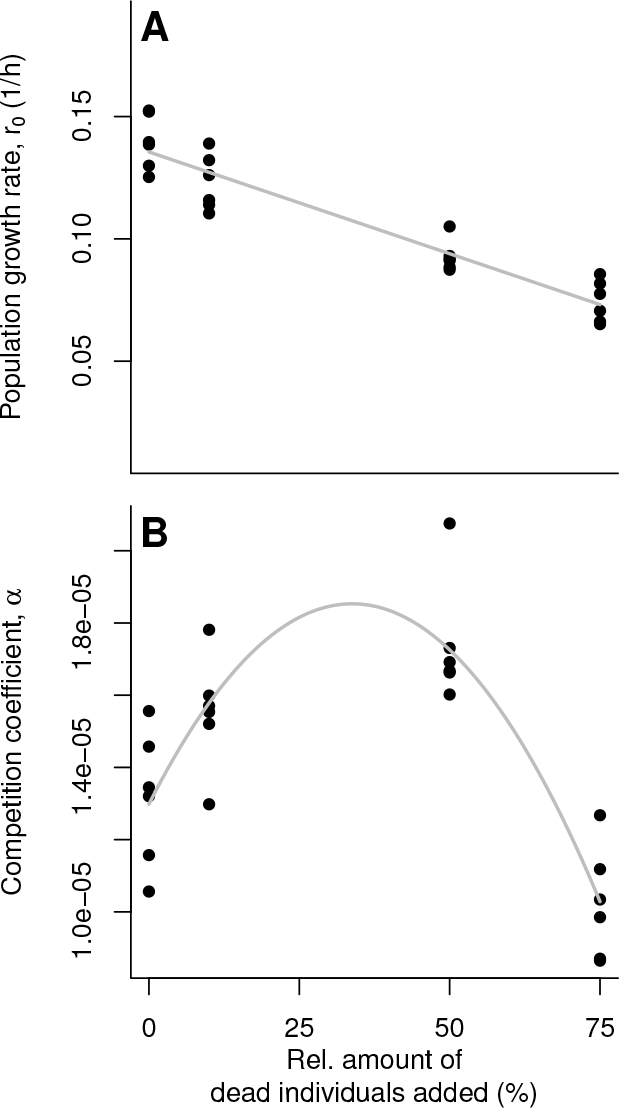
*Tetrahymena pyriformis* — chemical cues of dead conspecifics impact population growth and intraspecific competition. (A) We observed a negative effect of chemical cues on growth rates (*r*_0_; LM: *N* = 24(6), *t* = −12.79, *df* = 22, *p* < 0.001) and (B) a quadratic effect on the competition coefficient (*α*; LM, linear term: *N* = 24(6), *df* = 21, *t* = 6.74, *p* < 0.001; quadratic term: *N* = 24(6), *df* = 21, *t* = −7.54, *p* < 0.001). Movement behaviour did not differ significantly between the treatments, however a weakly significant effect of time was observed as excepted (Fronhofer et al., 2015) (GLM, treatment: *N* = 192, *df* = 189, *t* = 1.14, *p* = 0.26; time: *N* = 192, *df* = 189, *t* = 2.09, *p* = 0.038, error family: Gamma). The potential consequences of these effects on range dynamics are analysed theoretically in Fig. S5.

**Supplementary Figure S3:**
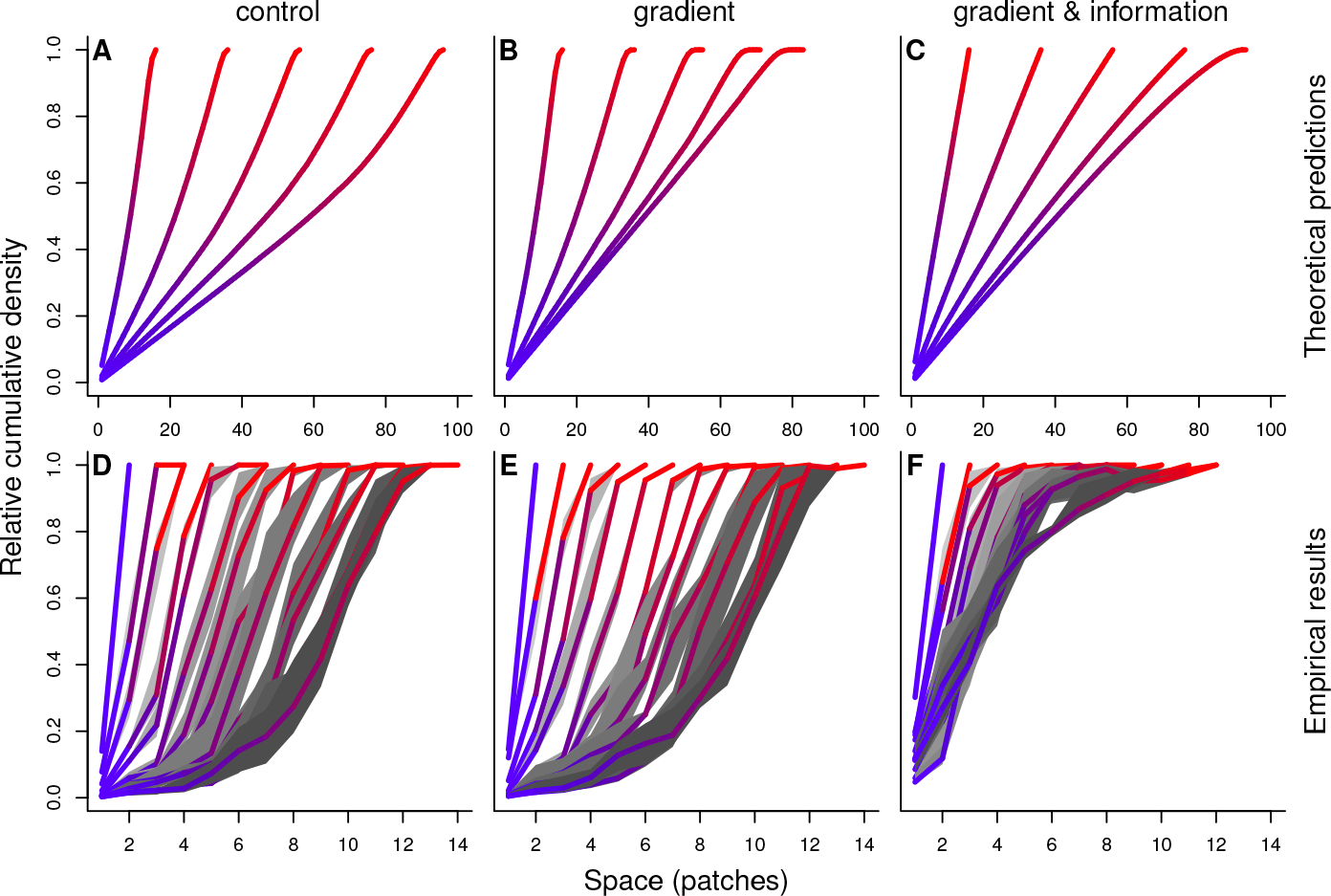
Cumulative density distributions of population densities over space during the range expansion — theoretical predictions and empirical results. Clearly, the cumulative density distributions in (A, control) and (B, gradient) are convex while the function is concave in the informed scenario (C). Note that after a sufficient amount of time the distributions in scenarios 2 and 3 should converge. Temporal snapshots: *t* = {10,30,50,70,90}. Parameter settings: λ_0_ = 14, *α*_0_ = 0.00001, *ρ* = 2, τ = 2. For the simulation results, we report medians over 20 replicate simulations (solid line; blue (range core) to red (range front)) and the 25th and 75th percentiles (grey shading; darker with time). As theoretically predicted (A-C), we find convex relationships in (D) and (E) while the relationship in concave in (F). Note that the differences between the scenarios are more pronounced than theoretically predicted. While this is not per se disturbing, since the model predictions are not thought to be quantitative, the differences can be explained by a shift in trait correlation structures (see main text and Fig. 3). For a statistical comparison of the density distributions see Fig. S4. For the empirical results, we report medians over 6 experimental replicates and the 25th and 75th percentiles.

**Supplementary Figure S4:**
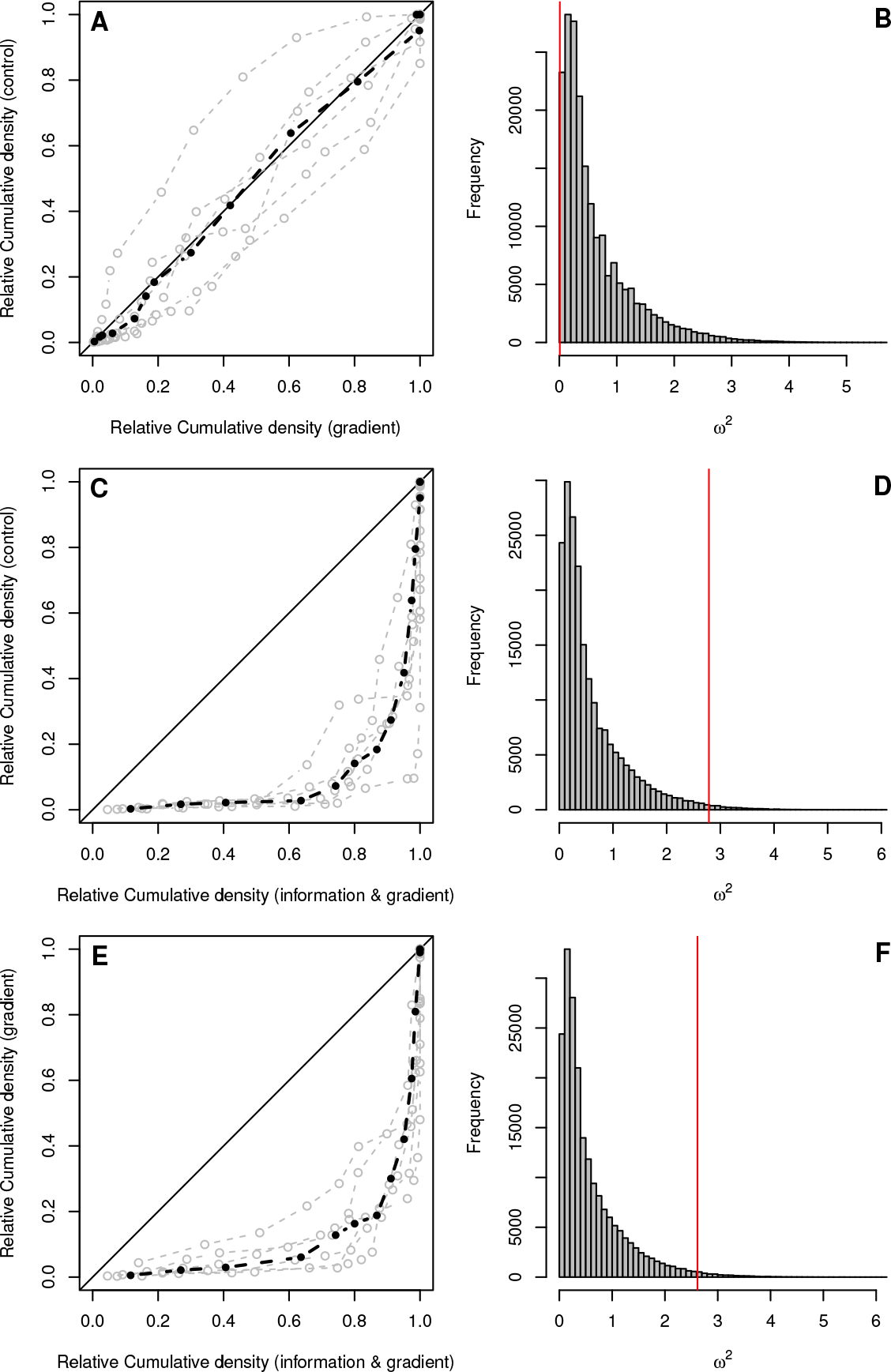
Statistical analysis of the empirical cumulative density distributions of population densities over space during the range expansion. (A, C, E) Probability-Probability plots. In case of identical distributions the data should lie on the diagonal (solid line). We here show the individual replicates (grey) and the median (black). The latter was used for the statistical analysis. (B, D, F) Distributions of the Cramer-von Mises (CvM) statistic (*ω*^2^) generated by resampling (200,000 draws) and the computed CvM statistic for the specific comparison (red line). CvM, control-gradient: *N* = 14, *ω*^2^ = 0.009, *p* = 0.99; CvM, control-information & gradient: *N* = 14, *ω*^2^ = 2.79, *p* = 0.024; CvM, gradient-information & gradient: *N* = 14, *ω*^2^ = 2.62, *p* = 0.028

**Supplementary Figure S5:**
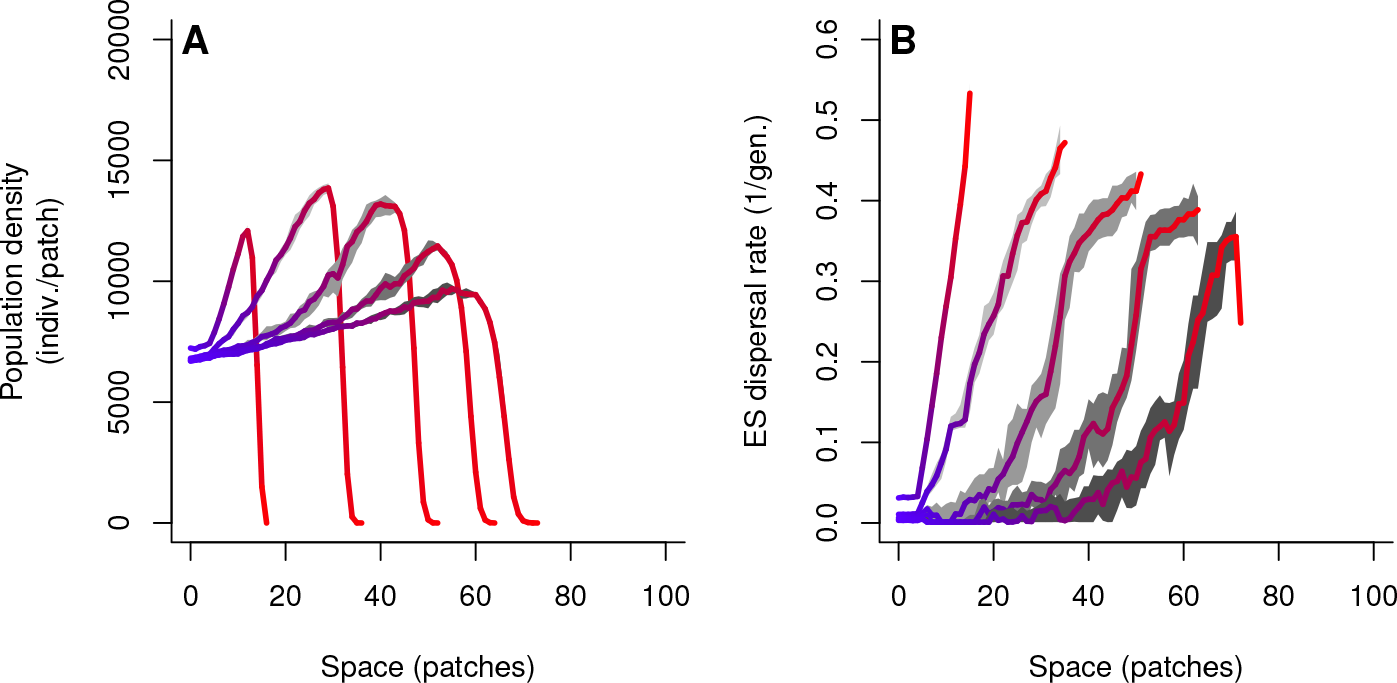
Predicted range dynamics into a combined mortality and decreasing fecundity gradient without information use. This scenario represents a sensitivity analysis for the direct negative effect of chemical cues on population growth as suggested by Fig. S2 A. With this analysis we intend to rule out the possibility that the experimental results of scenario 3 (“gradient & information”) are an artifact due to the negative side effects of chemical cues provided by dead conspecifics (Fig. S2) and not a result of information use. We therefore implemented a negative fecundity gradient that follows exactly the slope of the effect of chemical cues on growth rate reported in Fig. S2 A. The competition coefficient therefore also decreases following Eq. 3. As one can clearly see, this combined gradient does not destroy the pattern of increasing densities towards the range front (convex cumulative population density distribution). Consequently, we are confident that, while the negative effects of chemical cues exist and certainly act in our experiments, the density and movement patterns we report are not mere artifacts linked to these concurrent effects.

**Supplementary Figure S6:**
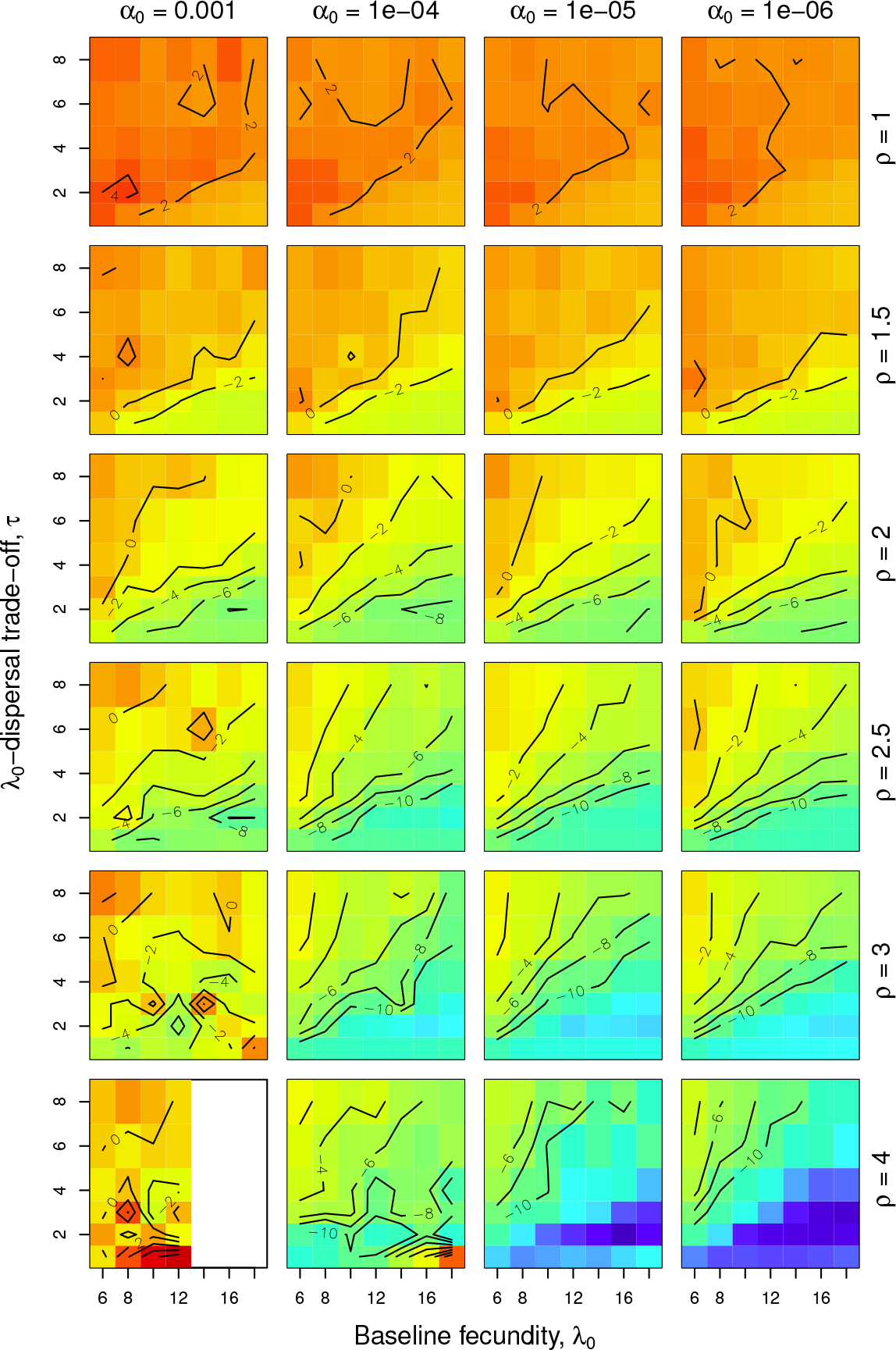
Sensitivity analysis: control. Convexity (concavity) of the relative cumulative population density distribution. The index reported here is the difference between a focal cumulative density distribution function and a linear connection between its first and last point. Negative (positive) values in blue (red) indicate convex (concave) cumulative population density distributions, i.e., higher (lower) densities at range fronts compared to range cores. All analyses were conducted on the median distribution of population densities over all 20 replicates from the end of a simulation run (*t* = 95).

**Supplementary Figure S7:**
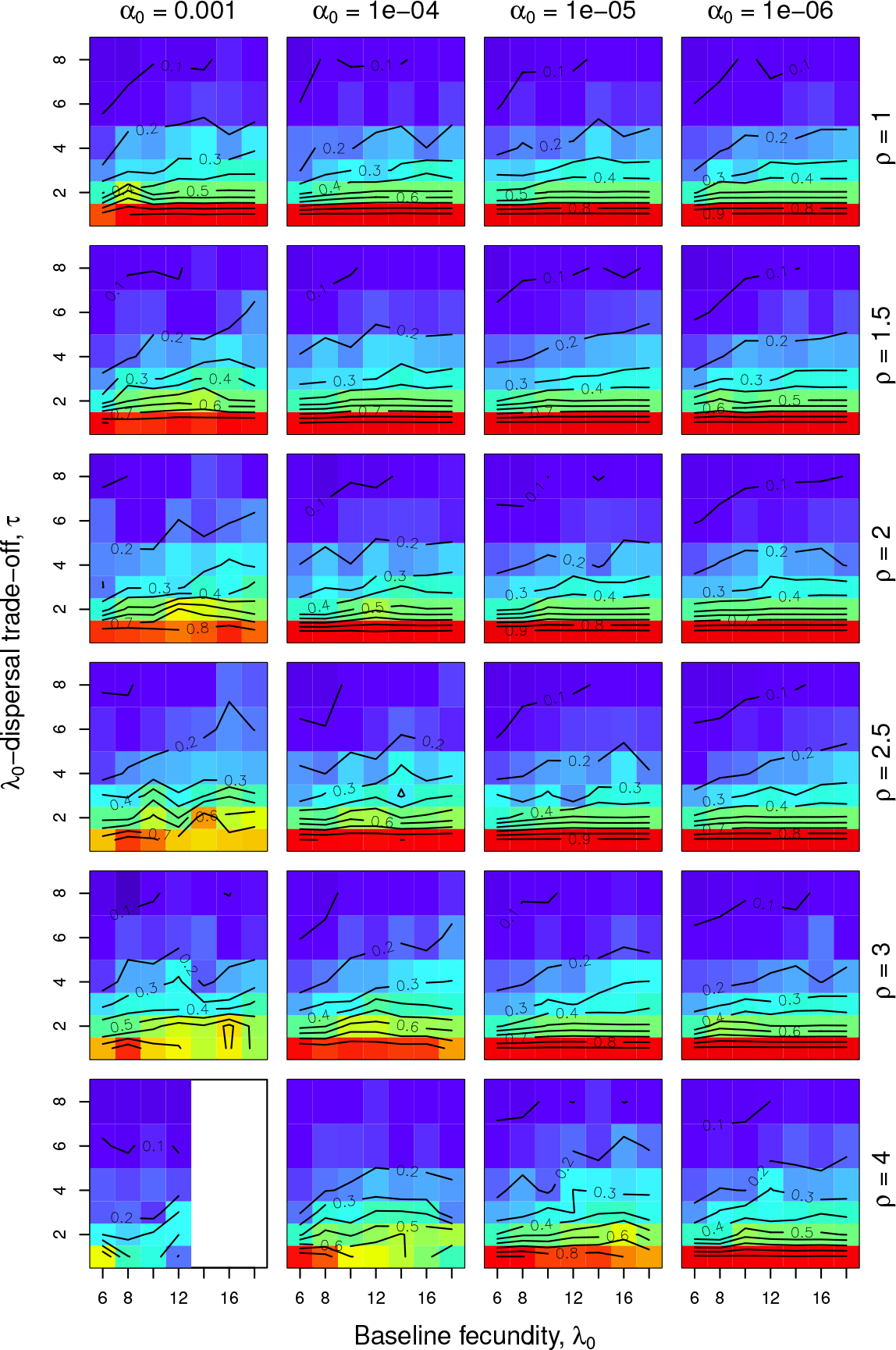
Sensitivity analysis: control. Difference between evolutionarily stable dispersal strategies in range range cores and range fronts. All analyses were conducted on the median spatial profile of evolutionary stable dispersal strategies over all 20 replicates from the end of a simulation run (*t* = 95).

**Supplementary Figure S8:**
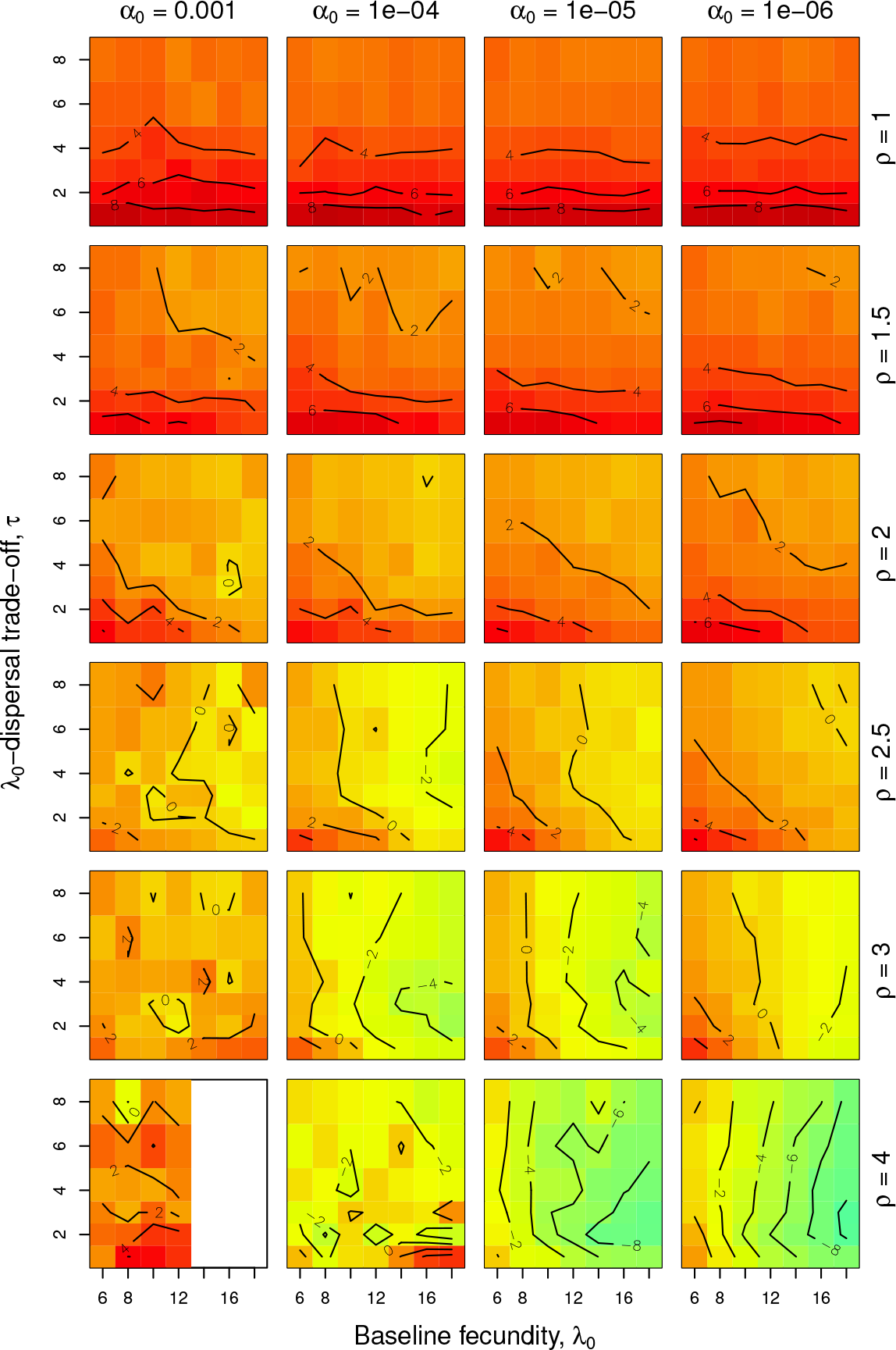
Sensitivity analysis: gradient. Convexity (concavity) of the relative cumulative population density distribution. The index reported here is the difference between a focal cumulative density distribution function and a linear connection between its first and last point. Negative (positive) values in blue (red) indicate convex (concave) cumulative population density distributions, i.e., higher (lower) densities at range fronts compared to range cores. All analyses were conducted on the median distribution of population densities over all 20 replicates from the end of a simulation run (*t* = 95).

**Supplementary Figure S9:**
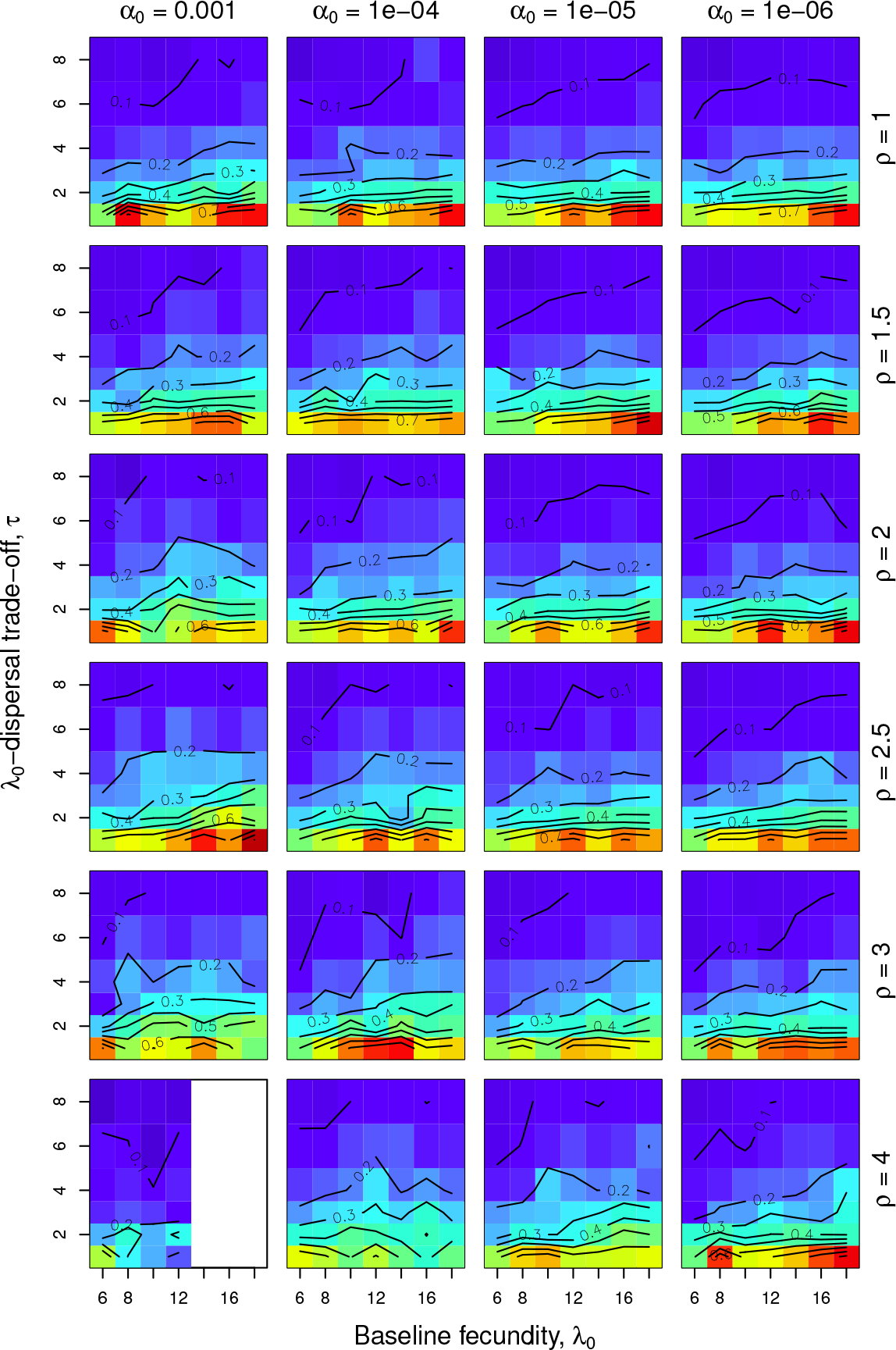
Sensitivity analysis: gradient. Difference between evolutionarily stable dispersal strategies in range range cores and range fronts. All analyses were conducted on the median spatial profile of evolutionary stable dispersal strategies over all 20 replicates from the end of a simulation run (*t* = 95).

**Supplementary Figure S10:**
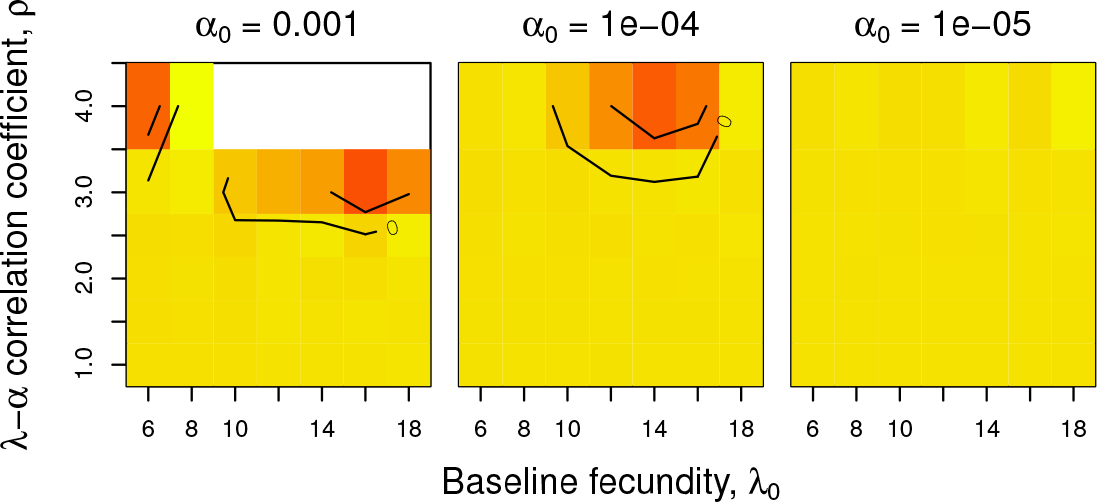
Sensitivity analysis: gradient & information. Convexity (concavity) of the relative cumulative population density distribution. The index reported here is the difference between a focal cumulative density distribution function and a linear connection between its first and last point. Negative (positive) values in blue (red) indicate convex (concave) cumulative population density distributions, i.e., higher (lower) densities at range fronts compared to range cores. All analyses were conducted on the median distribution of population densities over all replicates (between 1 and 10) from the end of a simulation run (*t* = 95).

## Supplementary tables

**Supplementary Table S1:** Important parameters of the evolutionary individual-based model, their meaning and tested values. Standard values are underlined.

| Parameter | Values | Meaning |
| --- | --- | --- |
| $\lambda_0$ | 6, 8, 10, 12, <u>14</u>, 16, 18 | baseline fecundity |
| $\alpha_0$ | 1E-03, 1E-04, <u>1E-05</u>, 1E-06 | baseline competition coefficient |
| $\rho$ | 1, 1.5, <u>2</u>, 2.5, 3, 4 | λ-α correlation coefficient |
| $\tau$ | 1, <u>2</u>, 3, 4, 6, 8 | λ-dispersal trade-off strength |

## References

Hill, J.K., Thomas, C.D. & Huntley, B. 1999 Climate and habitat availability determine 20th century changes in a butterflys range margin. Proc. R. Soc. B-Biol. Sci. 266, 1197–1206. doi:10.1098/rspb.1999.0763.

Parmesan, C., Ryrholm, N., Stefanescu, C., Hill, J.K., Thomas, C.D., Descimon, H., Huntley, B., Kaila, L., Kullberg, J. et al., 1999 Poleward shifts in geographical ranges of butterfly species associated with regional warming. Nature 399, 579–583. doi:10.1038/21181.

Parmesan, C. & Yohe, G. 2003 A globally coherent fingerprint of climate change impacts across natural systems. Nature 421, 37–42. doi:10.1038/nature01286.

Kelly, A.E. & Goulden, M.L. 2008 Rapid shifts in plant distribution with recent climate change. Proc. Natl. Acad. Sci. U. S. A. 105, 11823–11826. doi:10.1073/pnas.0802891105.

Hastings, A., Cuddington, K., Davies, K.F., Dugaw, C.J., Elmendorf, S., Freestone, A., Harrison, S., Holland, M., Lambrinos, J. et al. 2005 The spatial spread of invasions: new developments in theory and evidence. Ecol. Lett. 8, 91–101. doi:10.1111/j.1461-0248.2004.00687.x.

Holt, R.D., Keitt, T.H., Lewis, M.A., Maurer, B.A. & Taper, M.L. 2005 Theoretical models of species’; borders: single species approaches. Oikos 108, 18–27.

Burton, O.J., Pillips, B.L. & Travis, J.M. J. 2010 Trade-offs and the evolution of life-histories during range expansion. Ecol. Lett. 13, 1210–1220.

Dytham, C. 2009 Evolved dispersal strategies at range margins. Proc. R. Soc. B-Biol. Sci. 276, 14071413.

Holt, R.D. & Barfield, M. 2011 Theoretical perspectives on the statics and dynamics of species’; borders in patchy environments. Am. Nat. 178, S6–S25.

Perkins, A.T., Phillips, B.L., Baskett, M.L. & Hastings, A. 2013 Evolution of dispersal and life history interact to drive accelerating spread of an invasive species. Ecol. Lett. 16, 1079–1087. doi:10.1111/ele.12136.

Melbourne, B.A. & Hastings, A. 2009 Highly variable spread rates in replicated biological invasions: fundamental limits to predictability. Science 325, 1536–1539.

Giometto, A., Rinaldo, A., Carrara, F. & Altermatt, F. 2014 Emerging predictable features of replicated biological invasion fronts. Proc. Natl. Acad. Sci. U. S. A. 111, 297–301. doi:10.1073/pnas.1321167110.

Fronhofer, E.A. & Altermatt, F. 2015 Eco-evolutionary feedbacks during experimental range expansions. Nat. Commun. 6, 6844. doi:10.1038/ncomms7844.

Kubisch, A., Hovestadt, T. & Poethke, H.J. 2010 On the elasticity of range limits during periods of expansion. Ecology 91, 3094–3099.

Kubisch, A., Holt, R.D., Poethke, H.J. & Fronhofer, E.A. 2014 Where am I and why? Synthesising range biology and the eco-evolutionary dynamics of dispersal. Oikos 123, 5–22. doi:10.1111/j.1600-0706.2013.00706.x.

Louthan, A.M., Doak, D.F. & Angert, A.L. 2015 Where and when do species interactions set range limits? Trends Ecol. Evol. 30, 780792. doi:10.1016/j.tree.2015.09.011.

Clobert, J., Le Galliard, J.F., Cote, J., Meylan, S. & Massot, M. 2009 Informed dispersal, heterogeneity in animal dispersal syndromes and the dynamics of spatially structured populations. Ecol. Lett. 12, 197–209.

Altermatt, F., Fronhofer, E.A., Garnier, A., Giometto, A., Hammes, F., Klecka, J., Legrand, D., Machler, E., Massie, T.M. et al. 2015 Big answers from small worlds: a user’;s guide for protist microcosms as a model system in ecology and evolution. Methods Ecol. Evol. 6, 218–231. doi:10.1111/2041-210X.12312. The detailed protocols (that appear in the supplement) are available at http://emeh-protocols.readthedocs.org/.

Phillips, B.L., Brown, G.P., Webb, J.K. & Shine, R. 2006 Invasion and the evolution of speed in toads. Nature 439, 803–803. doi:10.1038/439803a.

Bonte, D., Van Dyck, H., Bullock, J.M., Coulon, A., Delgado, M., Gibbs, M., Lehouck, V., Matthysen, E., Mustin, K. et al. 2012 Costs of dispersal. Biol. Rev. 87, 290–312. doi:10.1111/j.1469-185X.2011.00201.x.

Matessi, C. & Gatto, M. 1984 Does k-selection imply prudent predation? Theor. Popul. Biol. 25, 347–363. doi:10.1016/0040-5809(84)90014-5.

Beverton, R. J. H. & Holt, S.J. 1957 On the dynamics of exploited fish populations. London: Chapman & Hall.

Hamilton, W.D. & May, R.M. 1977 Dispersal in stable habitats. Nature 269, 578–581. doi:10.1038/269578a0.

Kubisch, A., Fronhofer, E.A., Poethke, H.J. & Hovestadt, T. 2013 Kin competition as a major driving force for invasions. Am. Nat. 181, 700–706. doi:10.1086/670008.

Phillips, B.L., Brown, G.P. & Shine, R. 2010 Life-history evolution in range-shifting populations. Ecology 91, 1617–1627. doi:10.1890/09-0910.1.

Shine, R., Brown, G.P. & Phillips, B.L. 2011 An evolutionary process that assembles phenotypes through space rather than through time. Proc. Natl. Acad. Sci. U. S. A. 108, 5708–5711. doi:10.1073/pnas.1018989108.

Cadotte, M.W. 2007 Competition-colonization trade-offs and disturbance effects at multiple scales. Ecology 88, 823–829. doi:10.1890/06-1117.

Pennekamp, F. & Schtickzelle, N. 2013 Implementing image analysis in laboratory-based experimental systems for ecology and evolution: a hands-on guide. Methods Ecol. Evol. 4, 483–492. doi:10.1111/2041-210X.12036.

Pennekamp, F., Schtickzelle, N. & Petchey, O.L. 2015 Bemovi, software for extracting behavior and morphology from videos, illustrated with analyses of microbes. Ecol. Evol. 5, 2584–2595. doi:10.1002/ece3.1529.

Fronhofer, E.A., Kropf, T. & Altermatt, F. 2015a Density-dependent movement and the consequences of the allee effect in the model organism Tetrahymena. J. Anim. Ecol. 84, 712–722. doi:10.1111/1365-2656.12315.

Fronhofer, E.A., Klecka, J., Melian, C. & Altermatt, F. 2015b Condition-dependent movement and dispersal in experimental metacommunities. Ecol. Lett. 18, 954–963. doi:10.1111/ele.12475. As a preprint on BioRxiv http://dx.doi.org/10.1101/017954.

Sbalzarini, I. & Koumoutsakos, P. 2005 Feature point tracking and trajectory analysis for video imaging in cell biology. J. Struct. Biol. 151, 182–195.

## References

Mallet, J. 2012 The struggle for existence: how the notion of carrying capacity, K, obscures the links between demography, darwinian evolution, and speciation. Evol. Ecol. Res. 14, 627–665.

Reznick, D., Bryant, M.J. & Bashey, F. 2002 r-and K-selection revisited: the role of population regulation in life-history evolution. Ecology 83, 1509–1520. doi:10.1890/0012-9658(2002)083[1509:raksrt]2.0.co;2.

Rueffler, C., Egas, M. & Metz, J. A. J. 2006 Evolutionary predictions should be based on individual-level traits. Am. Nat. 168, E148–E162. doi:10.1086/508618.

MacArthur, R. 1970 Species packing and competitive equilibrium for many species. Theor. Popul. Biol. 1, 1–11.

Schoener, T.W. 1973 Population growth regulated by intraspecific competition for energy or time: Some simple representations. Theor. Popul. Biol. 4, 56–84. doi:10.1016/0040-5809(73)90006-3.

Abrams, P. 2009 Determining the functional form of density dependence: Deductive approaches for consumer-resource systems having a single resource. Am. Nat. 174, 321–330. doi:10.1086/603627.

Verhulst, P.-F. 1838 Notice sur la loi que la population suit dans son accroissement. Correspondance Mathematique et Physique 10, 113–121.

Fronhofer, E.A., Kropf, T. & Altermatt, F. 2015 Density-dependent movement and the consequences of the allee effect in the model organism Tetrahymena. J. Anim. Ecol. 84, 712–722. doi:10.1111/1365-2656.12315.

